# Three-dimensional reconstructions of the internal structures of haustoria in parasitic Orobanchaceae

**DOI:** 10.1101/2020.05.07.083055

**Authors:** Natsumi Masumoto, Yuki Suzuki, Songkui Cui, Mayumi Wakazaki, Mayuko Sato, Kie Kumaishi, Arisa Shibata, Kaori M. Furuta, Yasunori Ichihashi, Ken Shirasu, Kiminori Toyooka, Yoshinobu Sato, Satoko Yoshida

**Author notes:** Graduate School of Medicine, Osaka University, Osaka, Japan (Yuki Suzuki). Authors for contact: Yoshida Satoko,; Songkui Cui. **Author contributions:** SC and SY conceived the idea of this study, designed the experiment, analyzed the data and wrote the manuscript. NM was responsible for color coding, FE-SEM and 3-D reconstruction. YSu and YSa developed methods to align section images, automated the color coding process, and provided crucial technical assistance for 3-D reconstruction. YI, AS and KK assisted with image handling. KT, MW and MS prepared serial thin sections and performed FE-SEM. KF analyzed the At*PEAR* promoter. KS provided critical comments on the manuscript. **Authors for contact details:** Satoko Yoshida, Division of Biological Science, Graduate School of Science and Technology, Nara Institute of Science and Technology, Ikoma, Nara, Japan, Songkui Cui, Institute for Research Initiatives, Division for Research Strategy, Nara Institute of Science and Technology, Ikoma, Nara, Japan.

## Abstract

Parasitic plants infect other plants by forming haustoria, specialized multicellular organs consisting of several cell types each of which has unique morphological features and physiological roles associated with parasitism. Understanding the spatial organization of cell types is, therefore, of great importance in elucidating the functions of haustoria. Here, we report a three-dimensional (3-D) reconstruction of haustoria from two Orobanchaceae species, the obligate parasite *Striga hermonthica* infecting rice and the facultative parasite *Phtheirospermum japonicum* infecting *Arabidopsis*. Our images reveal the spatial arrangements of multiple cell types inside haustoria and their interaction with host roots. The 3-D internal structures of haustoria highlight differences between the two parasites, particularly at the xylem connection site with the host. Our study provides structural insights into how organs interact between hosts and parasitic plants.

**One-sentence summary:** Three-dimensional image reconstruction was used to visualize the spatial organization of cell types in the haustoria of parasitic plants with special reference to their interaction with host roots.

## Introduction

Parasitic plants are higher plants that invade host plant tissues to acquire water and nutrients from their hosts. Plant parasitism independently evolved at least 12 times (Bennett and Mathews, 2006). Approximately 4,500 species in 28 families, representing 1% of all angiosperm species, are proposed to be parasitic plants (Westwood et al., 2010; Heide-Jørgensen, 2013). The common feature of parasitic plants is the ability to form a specialized invasive organ called a haustorium. Unlike the unicellular haustoria found in plant-infecting pathogenic fungi, haustoria in parasitic plants are multicellular organs that function in host attachment, host tissue invasion and establishing a vascular connection between the parasite and host for material transfer (Yoshida et al., 2016). Haustoria facilitate water and nutrient acquisition as well as translocation of RNA molecules, peptides and plant hormones between the host and parasite (Kim et al., 2014; Spallek et al., 2017; Shahid et al., 2018; Liu et al., 2019; Yoshida et al., 2019). Thus, haustoria act as efficient biological channels for interspecies material transfer, but the internal structures and physiological functions of haustoria are largely unexplored.

The Orobanchaceae family contains the largest number of root parasitic species and present various degrees of host dependency. An exception among Orobanchaceae family members is in the genus *Lindenbergia* of which all species are non-parasitic plants (Hjertson, 1995). Facultative parasites, including those in the genera *Triphysaria*, *Phtheirospermum*, and *Rhinanthus*, are able to live as autotrophs and reproduce seeds without a host, but will parasitize hosts when available. Obligate parasitic plants, including those in the genera *Striga*, *Orobanche*, and *Phelipanche*, cannot complete their life cycles in nature without a host. Some of the obligate parasites in the Orobanchaceae represent major agricultural threats because they parasitize important staple crops. For example, *Striga hermonthica*, commonly known as witchweed, infests rice, maize, sorghum, sugarcane and millet, affects millions of smallholder farmers in sub-Saharan Africa and, thus, has a large impact on global food security (Spallek et al., 2013; Runo and Kuria, 2018; Mutuku and Shirasu, 2019).

Haustorial morphology varies depending on the species. Facultative parasitic plants form lateral haustoria that develop on the sides of roots without permanently stopping meristematic activity of the roots. In contrast, obligate parasitic plants form terminal haustoria at the radicle tips by deforming the root apical meristem (Westwood et al., 2010). These obligate parasites have small seeds with limited nutrient reserves (Joel et al., 2012); thus, nutrient acquisition through terminal haustoria is crucial for the survival of obligate parasites. Some obligate parasites, such as *Striga*, are also able to form lateral haustoria on their secondarily emerged adventitious roots after successful establishment of a host connection via a terminal haustorium.

Despite the evidence for distinct haustorium types, the developmental processes and major internal cell structures are similar between lateral and terminal haustoria in the Orobanchaceae. The initial formation of a haustorium is provoked by host-derived small compounds, collectively called haustorium-inducing factors (HIFs) (Goyet et al., 2019). The HIF signals induce cell division and cell deformation as well as the extension of epidermal cells on the haustorium-forming site, called haustorial hairs (or papillae), that function in host attachment (Cui et al., 2016). Upon host contact, haustoria develop intrusive cells at their apex, acting at the front line of host penetration. The intrusive cells penetrate host tissues toward the host’s stele and, after successful entry into the host’s vascular system, parts of the intrusive cells differentiate into tracheary elements. In parallel, a mass of tracheary elements, called plate xylem, develop in the basal region of the haustorium (Heidejorgensen and Kuijt, 1995). These two types of xylem tissues are eventually connected in the middle of the haustorium to form a xylem bridge that facilitates molecular flow between the parasite and its host (Ishida et al., 2016; Wakatake et al., 2018). In the middle of a haustorium surrounding the xylem bridge, small cells have been observed (Wakatake et al., 2018; Goyet et al., 2019). In the facultative parasitic plant *Phtheirospermum japonicum*, procambium marker genes, such as homologs of *Arabidopsis HB15a*, *HB18* and *WOX4*, were highly expressed in these small cells; therefore, these cells were called procambium-like cells (Wakatake et al., 2018). After a host-parasite vascular connection is established, a cluster of parenchyma cells, called the hyaline body, are observed in many species including the obligate parasites *Striga* and *Alectra,* as well as the facultative parasite *Rhinanthus.* Hyaline cells forming the hyaline body have dense cytoplasm and are enriched with organelles having extracellular depositions (Visser et al., 1984; Pielach et al., 2014). Because hyaline bodies appear only in compatible host-parasite interactions, their roles have been postulated as being important for later stages of parasitic plant growth (Gurney et al., 2006; Pielach et al., 2014).

The structure of the vascular connection between hosts and parasites varies depending on the species (Smith et al., 2013). An obligate parasite *Striga hermonthica* and a facultative parasite *Rhinanthus minor* were reported to form oscula, tubular structures that directly penetrate host xylem vessels to form a conduit connection between the host and parasite (Dorr, 1997; Cameron et al., 2006); whereas, *Orobanche crenata* was reported to connect to host xylems by open pits (Dörr and Kollmann, 1976). A phloem connection has been reported in the holo (non-photosynthetic) parasitic species *Orobanche crenata* and *cumana* (Dorr and Kollmann, 1995; Krupp et al., 2019). No phloem cells were detected in the haustoria of host contact sites in hemi (photosynthetic) parasites including the obligate *Striga* and *Alectra* spp., as well as the facultative *Triphysaria* (Dorr et al., 1979; Ba, 1988; Heidejorgensen and Kuijt, 1993; Dorr, 1997; Kokla and Melnyk, 2018).

Although many histological studies and electron micrographs are available documenting host and parasitic plant interaction, the spatial relationship between host and parasite cells is not well understood. To overcome this issue, we have developed complete three-dimensional images of haustoria. Although fluorescence images taken by confocal microscopy are widely used for reconstructing 3-D images, this approach is challenging to use for haustoria due to their thick tissues. In addition, haustorial cell type-specific gene markers have not yet been fully identified in parasitic plants. Thus, we constructed 3-D images of haustoria from serial thin sections of *S. hermonthica* and *P. japonicum* and visualized individual cell types inside haustoria that were interacting with their hosts.

## Results

### 3-D reconstruction of an *S. hermonthica* haustorium infecting a rice root

To reconstruct a 3-D image of a haustorium, we followed the workflow scheme shown in Fig. 1. In brief, the protocol included: 1) tissue preparation, sectioning, staining and taking photographs, 2) aligning photographed sections, 3) manual painting of each cell type in every fifth image, 4) automated painting based on manually painted images and cellular segmentation, and 5) stacking of the images and reconstruction of a 3-D structure. For this purpose, we prepared serial thin sections from a resin-embedded *S. hermonthica* haustorium after infecting rice for two weeks (Fig. 1A, C). In total, 526 1 μM-thick sections were recovered, and among them 374 consecutive sections covering an entire haustorium were selected for further analysis. Each slide was stained with toluidine blue, and the image was photographed using a microscope. Because each section was manually placed on a glass slide, the positions and angles varied among the sections. Therefore, we applied intensity-based rigid image registration to align photographed sections using an image registration library (Marstal et al., 2016). In addition, some sections had wrinkles and unequal staining that caused non-rigid deformation. To avoid tissue discontinuity due to non-rigid deformation, we also applied intensity-based non-rigid image registration to the images having wrinkled sections (for details see Materials and Methods).

**Figure 1.**
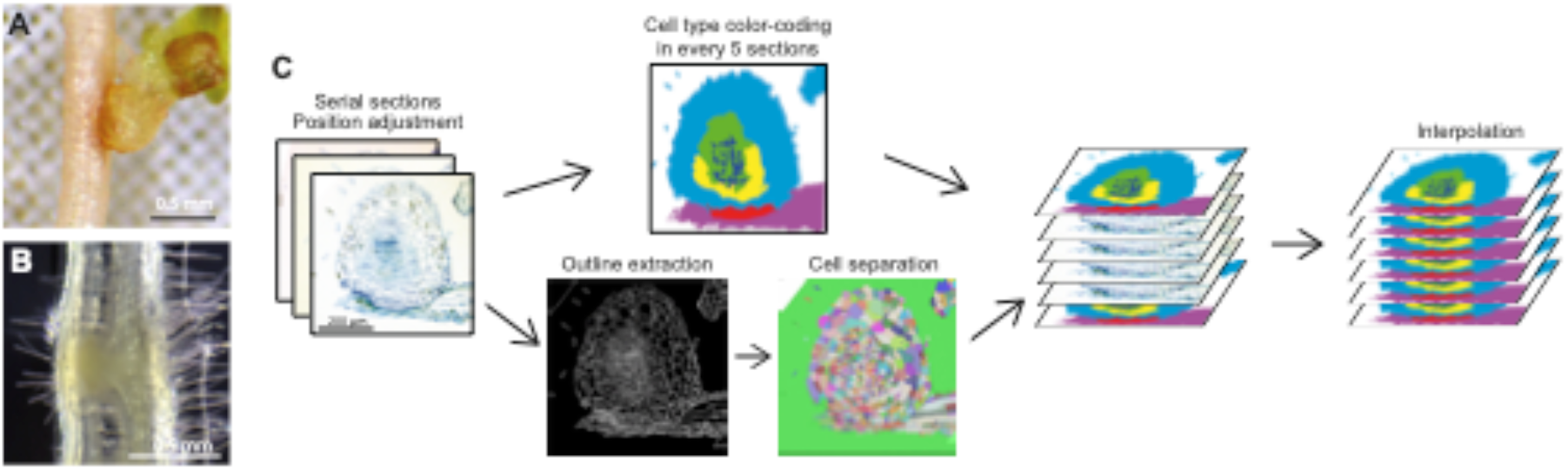
3-D reconstructions of *S. hermonthica* and *P. japonicum* haustoria. A, A terminal haustorium of *S. hermonthica* that infected a rice root for 2 weeks. B, A lateral haustorium of *P. japonicum* that infected an *Arabidopsis* root for 9 days. Scale bars: A, B = 500 μm, C = 200 μm. C, A workflow diagram of 3-D reconstruction using *S. hermonthica* section images.

### Defining cell types in a *S. hermonthica* haustorium

To visualize the structure of cell types, we used a color-coding system; each cell type was painted with a different color. We have categorized the parasite cell types into six categories, *i.e.* cortical cells, hyaline cells, procambium-like cells, intrusive cells, tracheary elements, and sieve elements. Host rice cells were categorized into two cell types, vascular cells including xylem, phloem and procambium, and other cells including endodermal, cortical and epidermal cells. The following morphological characteristics were used as criteria for color-coding different cell types, and each cell type was observed using field-emission scanning electron microscopy (FE-SEM) to confirm intracellular structures (Fig. 2). In section images, larger cells in the periphery of the haustorium and smaller cells in the center of the haustorium were observed. Larger cells at the haustorium’s periphery are cortical cells that have vacuolated intracellular structures (Fig. 2A, B, H, K). Smaller cells at the center of the haustorium contain five cell types including hyaline body, procambium-like cells, intrusive cells, tracheary elements and sieve elements (SE). (Fig. 2A). Hyaline cells, constituting the hyaline body, were stained light blue by toluidine blue. The extracellular spaces of hyaline cells were stained dark blue by toluidine blue and contained highly electron dense material as observed by FE-SEM, indicating extracellular deposits, a typical feature of hyaline bodies (Fig. 2B-D). FE-SEM analysis revealed that the cytosol of hyaline cells was densely occupied by irregular membrane structures, likely thickened smooth endoplasmic reticulum (ER) (Fig. 2D, E). Among the small, round cells, we categorized cells missing the features of hyaline cells as procambium-like cells, as observed previously in *P. japonicum* cellular structure (Wakatake et al., 2018). Procambium-like cells displayed a lighter cytosolic density with fewer ER structures and a tight membrane contact with each other compared to hyaline cells (Fig. 2B, C, F). Because procambium cells in *Striga* shoots are also small and morphologically indistinguishable from haustorial procambium-like cells on section images, we applied the same color to both cell types. Intrusive cells were characterized by their slender cell shape and palisade-like alignment against host vasculatures (Fig. 2H-J). Tracheary elements are dead cells with thick secondary cell walls, recognized by their empty lumen and pitted outline (Fig. 2F, I). Differentiating tracheary elements had only an empty lumen (Asterisk in Fig. 2I). Some tracheary elements directly penetrated host xylem vessels and formed structures previously described as oscula (Fig. 2H, L) (Dorr, 1997). Cell wall continuity between haustorial tracheary elements and host vessels was frequently observed at the host-parasite interface (Fig. 3A). SE showed less condensed cytoplasm and thickened cell walls. FE-SEM analysis detected characteristic plastid structures (Fig. 2C, G), called sieve-element plastids in SE cells (Behnke, 1991). We observed SE cells near the procambium-like cells in the basal region of the haustorium, but not in the central part of the haustorium (Fig. 2A, C). For the host, vascular tissues and the other cell types were marked with different colors. Finally, using the above criteria, we labeled cells in the aligned images. Since manually labeling cells in every section image is time consuming, we developed a semi-automated labeling method. In brief, color codes were manually painted on every fifth image, and the rest of the images were labeled by an automated interpolation algorithm (for details, see Materials and Methods). The resulting color images were manually checked and painting mistakes were corrected.

**Figure 2.**
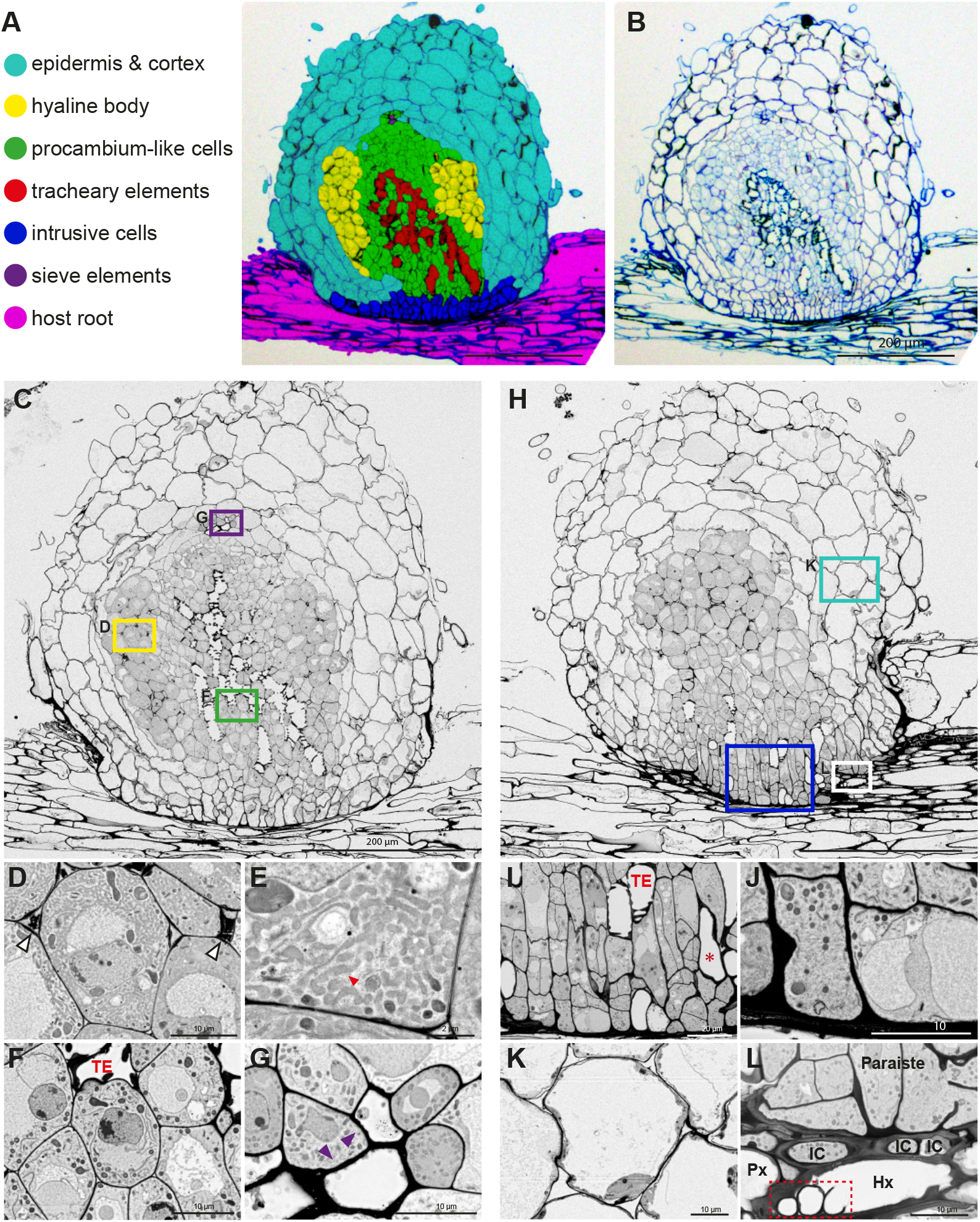
Tissue distribution and morphological characteristics of haustorial cells in *S. hermonthica*. A, Distribution of individual cell types in a *S. hermonthica* haustorium that infected rice for 2 weeks. B, Original toluidine blue-stained section used in A. C-G, FE-SEM images from a section in the center of a haustorium with a focus on the xylem bridge. Magnified images of hyaline cells (D, E), procambium-like cells (F) and sieve elements (purple arrowheads, G) from C are shown in the lower panels. E shows enrichment of smooth-ER like structures (red arrowhead) in a hyaline cell magnified from D. White arrowheads in D indicate the accumulation of electron dense extracellular materials in the hyaline body. H-L, FE-SEM images from a section on the side of a haustorium with a focus on intrusive cells. Below H are magnified images of intrusive cells (I, J), cortex cells (K) and oscula structures (red dashed box) within a host vessel element (L). L shows the host xylem (HX) surrounded by intrusive cells (IC) and a parasite tracheary element (PX). The corresponding positions of magnified images are indicated by colored boxes in C and H. TE: Tracheary element. Asterisk: differentiating TE. Scale bars, A-C, H = 200 μm; D, F, G, J-L = 10 μm; E = 2 μm; I = 20 μm.

**Figure 3.**
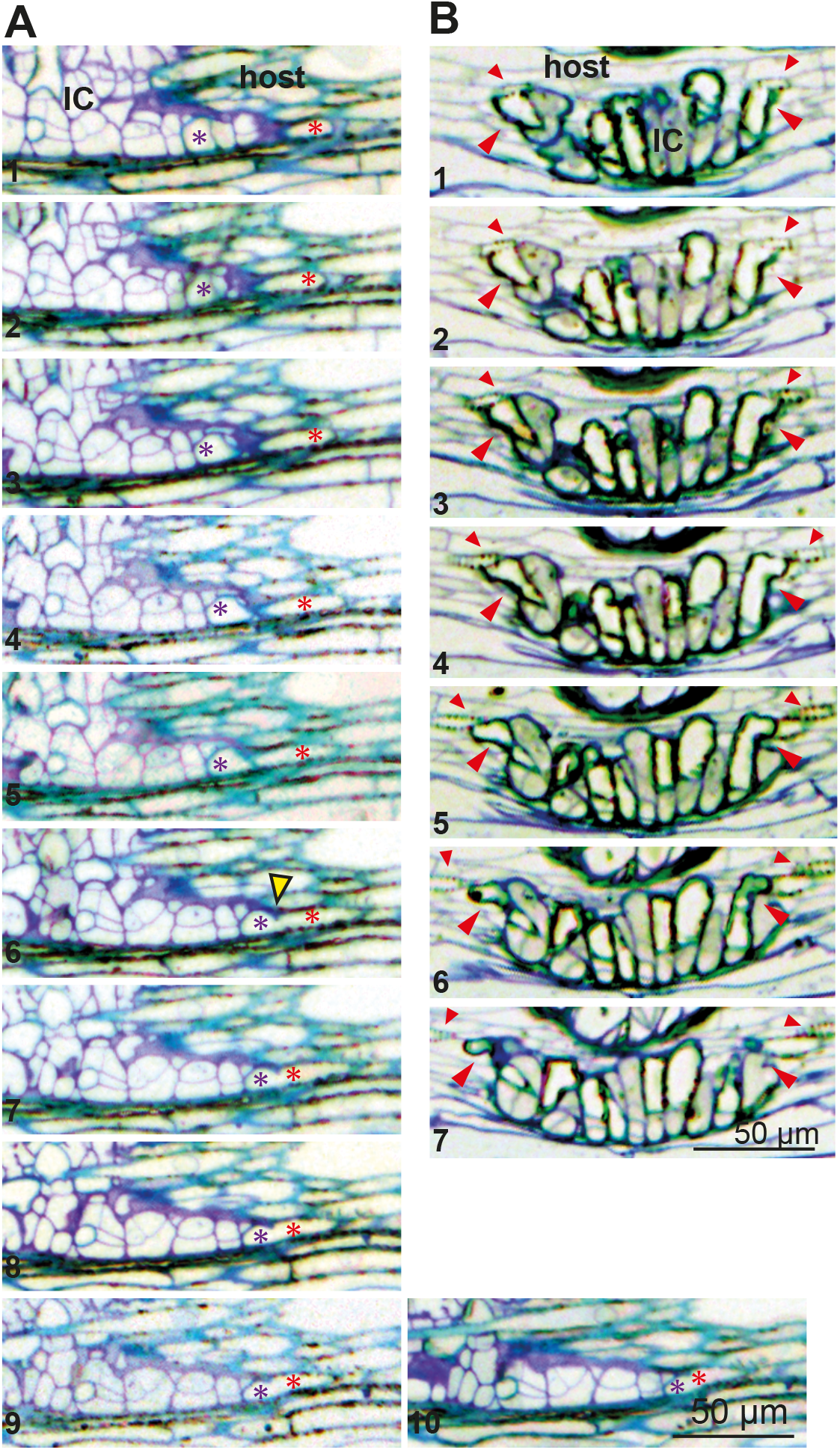
Interaction of parasite tracheary elements with host xylem at the interface. A, Continuous section images of the *S. hermonthica*-rice interface showing cell wall continuity (yellow arrowhead) between a tracheary element of *S. hermonthica* haustorium (purple asterisks) and a tracheary element of rice (red asterisks). **B,** Continuous section images of the *P. japonicum*-*Arabidopsis* interface showing the attachment of two *P. japonicum* tracheary elements (large arrowheads) next to host root xylem (small arrowheads). Note that no cell wall continuity was observed between the host xylem and parasite xylem. Scale bars, 50 μm.

### 3-D visualization of the internal structure of a *S. hermonthica* haustorium

All painted images were stacked to form a 3-D image and converted into polygon meshes that represent the surface structures of cell types. The polygon meshes were visualized by the open source program Paraview (Fig. 4, Movie 1-3). Opaque images visualized the surface structure of the haustorium (Supplemental Movie 1). Reconstruction of the tubular haustorial hairs indicates successful alignment of the serial sections (Fig. 4A). The semi-translucent 3-D image highlights the internal structure of a haustorium (Fig. 4B; Supplemental Movie 2). Xylem strands were continuous from the xylem vessels of a *Striga* shoot to the plate xylem and the xylem bridge in the haustorium (Fig. 4C-E; Supplemental Movie 3). Procambium-like cells and procambium cells surrounded the xylem strands (Fig. 4C, D; Supplemental Movie 3). Plate xylem formed a single aggregate at the base of the haustorium, and below that, ten xylem bridges seamlessly connected to the host xylem cells (Fig. 4E). Interestingly, interface xylem cells appeared to bend their tips outwards along the longitudinal axis of the host’s vasculature (Fig. 4E). A hyaline body assumed a ring shape surrounding procambium-like cells (Fig. 4C, D, F). Intrusive cells constituted the outermost layer of the haustorial apex region and covered most of interface area (Fig. 4B, D). Inside a *Striga* shoot, a few SE cell files extended in parallel with the shoot xylem. The tips of SE files were located above the xylem bridge in the vicinity of the plate xylem (Fig. 4C, D).

**Figure 4.**
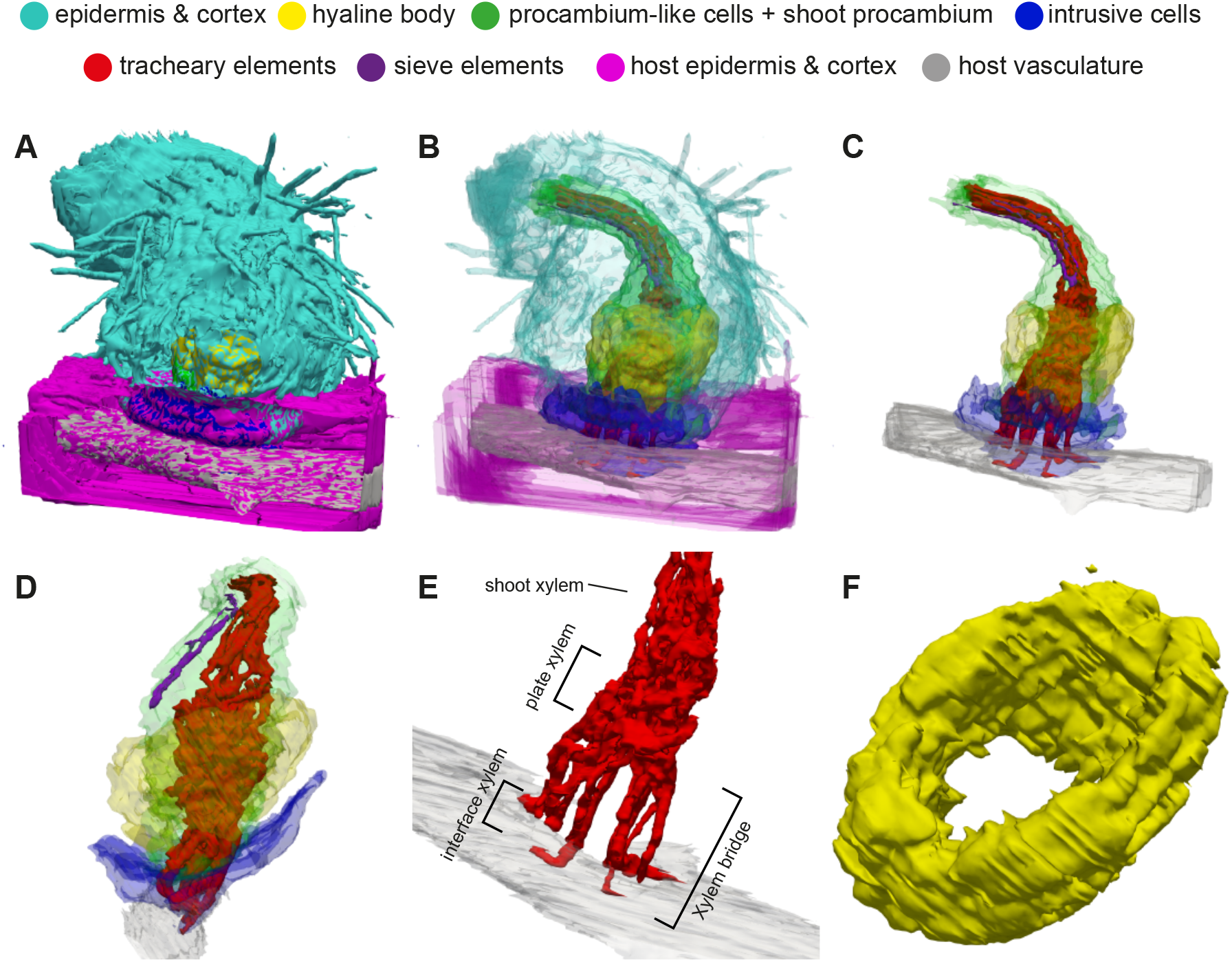
3-D images of a *S. hermonthica* haustorium. A, Surface structure. B, Semi-transparent view of the haustorium and host root. C, D, Internal haustorial cells and host root vasculature from the front (C) and side (D) views, respectively. E, A close-up view of the interface xylem, plate xylem and shoot xylem of *Striga* with host vasculature. F, Close up view of a hyaline body.

### Defining cell types in a *P. japonicum* haustorium infecting *Arabidopsis* roots

To reconstruct 3-D structures of a facultative parasitic plant, we infected *Arabidopsis* seedlings with *P. japonicum* (Fig. 1B). After 9 days, the tissues were fixed, embedded in resin and sectioned. We recovered 378 1 μm-thick serial sections from the *P. japonicum* haustorium (Fig. 5). Three-dimensional reconstruction was accomplished using the same procedure as that used for *Striga*. We defined cell types in the *P. japonicum* haustorium as follows. In the *P. japonicum* haustorium, hyaline cells were not observed. Instead, cells with a dense cytosol, enriched organelles and thickened primary cell walls were observed next to the xylem bridge (Fig. 5A-D). These cells have been also found in other facultative parasites including *Triphysaria*, *Rhinanthus*, *Odontites* and *Melampyrum* and were previously described as paratracheal parenchyma (PP) (Heidejorgensen and Kuijt, 1995; Pielach et al., 2014). Thus, we defined six cell types in the *P. japonicum* haustorium, i.e. cortical cells (Fig. 5E), PP (Fig. 5D), procambium-like cells (Fig. 5F-H), tracheary elements (Fig. 5D), intrusive cells (Fig. 5J, K) and SE (Fig. 5L, M). Procambium-like cells in *P. japonicum* were highly vacuolated and surrounded the PP and tracheary elements in the center (Fig. 5A-C). Detailed observations on the subcellular contents showed that these procambium-like cells can be divided into two groups (Fig. 5F). Those at the base and center of the haustorium (shown in red, Fig. 5F) were occupied by large vacuoles that contain many single- and multi-membrane small vesicles (Fig. 5G). Membrane invaginations were often observed adjacent to the vacuole membrane (arrowheads, Fig. 5G), indicating that the cytosolic matrix and organelles are actively degraded in these cells. In contrast, those located proximally to the host (blue color, Fig. 5F) had a relatively large cytosolic area that accumulated many atypical, unknown single-membrane structures (Fig. 5H). Unlike *S. hermonthica*, oscula structures were not observed at the invasion site (Fig. 3B). Cell wall continuity between the interface xylem and host vessel elements was not found at the interface (Fig. 3B, Fig. S2), indicating that intrusive cells of *P. japonicum* do not penetrate *Arabidopsis* xylem. SE were observed in the haustorium but only near the plate xylem (Fig. 5L, M). We could not determine whether these SE made continuity with root SE in the section images because of the difficulty in finding root SE in longitudinal sections.

**Figure 5.**
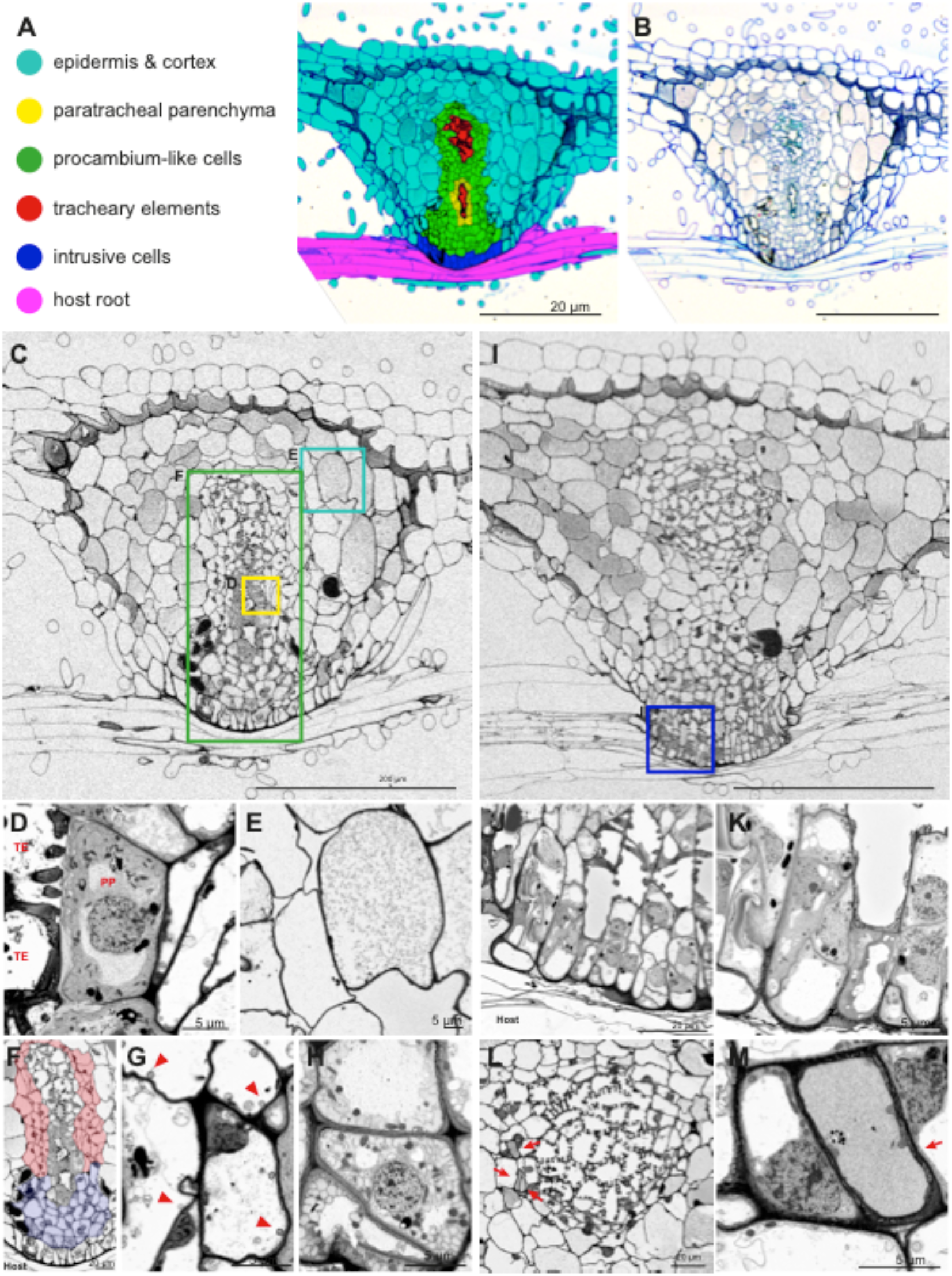
Tissue distribution and subcellular characteristics of individual cell types in a *P. japonicum* haustorium. A, Distribution of individual cell types in a haustorium that infected *Arabidopsis* for 9 days. B, Original toluidine blue-stained section image used in A. C-H, FE-SEM images from a section in the middle of a haustorium with the focus on a xylem bridge. Magnified images of paratracheal parenchyma (PP) and tracheary elements (TE) (D), cortex cells (E), procambium-like cells (F-H) are shown below C. Two types of procambium-like cells are indicated by red and blue (F), which are magnified in G and H, respectively. Arrowheads in G indicate vacuolar membrane invaginations. I-M, FE-SEM images at the side of a haustorium with a focus on intrusive cells. Magnified images of intrusive cells (J, K), plate xylem (L) and sieve elements (M) are shown in the panels below I. Arrowheads indicate sieve elements around the plate xylem. Scale bars, A, B, F, J, L = 20 μm; C, I = 200 μm; D, E, G, H, K, M: 5 μm; G: 10 μm.

### 3-D visualization of the internal structure of a *P. japonicum* haustorium

The 3-D model of a *P. japonicum* haustorium (Fig. 6, Supplemental Movie 4-6) revealed that the haustorial surface is densely covered with haustorial hairs (Fig. 6A). Analysis of the surface structure highlighted the structural differences between haustorial hairs and root hairs. Root hairs facing toward the host roots (haustorial hairs) are wavy and thicker with a higher density compared to the hairs located opposite the host roots (root hairs), which are thinner and straight (Supplemental Movie 4). Some haustorial hairs appeared to stick to the host surface and be firmly attached to the host root, supporting the notion that haustorial hairs are recruited for host attachment (Cui et al., 2016). Plate xylem was observed at the base of the haustorium near the *P. japonicum* root xylem, and a single-stranded xylem bridge connected to four interface xylem strands that interacted with the host xylem (Fig. 6D, E). Interestingly interface xylem cells do not have straight contact with host xylem but assumed an arc shape to wrap around the host xylem (Fig. 6D, E). The middle of the xylem bridge was encircled by PP cells, whereas the rest of the xylem bridge and the plate xylem were surrounded by procambium-like cells (Fig. 6C). Similar to *S. hermonthica*, intrusive cells of *P. japonicum* covered the host’s surface at the site where the haustorium first interacted with the host (Fig. 6F). Sieve elements in the haustorium were detected as two small clusters around the xylem plate at the outer boundary of procambium-like cells (Fig. 6G, H). Since we were not able to identify root SE in our experimental system, the continuity of haustorial SE with root SE could not be resolved in 3-D. To answer this question, we expressed an SE marker construct, *p*At*PEAR1::tdTomato-nls*, in which the *Arabidopsis PEAR1* promoter was fused with a *tdTomato* red fluorescent protein gene and a nuclear localization signal, in *P. japonicum* hairy roots (Fig. 6I, J). In *Arabidopsis*, *PEAR1* is known to be specifically expressed in protophloem sieve elements (Miyashima et al., 2019). In *P. japonicum* hairy roots, the fluorescence signal was observed in the longitudinal cell files within the root vasculature (Fig. 6I), indicating that this construct can serve as a SE marker in *P. japonicum*. A number of haustorial cells next to the plate xylem also expressed the marker gene, giving rise to a similar SE accumulation pattern to that observed in our 3-D image (Fig. 6C, I). This observation strongly supports the hypothesis that SE are present in the haustorium in the vicinity of the plate xylem. Notably, the fluorescence signal appeared to be continuous at the junction between the parasite’s root and the haustorium, indicating that haustorial SE are connected to root SE.

**Figure 6.**
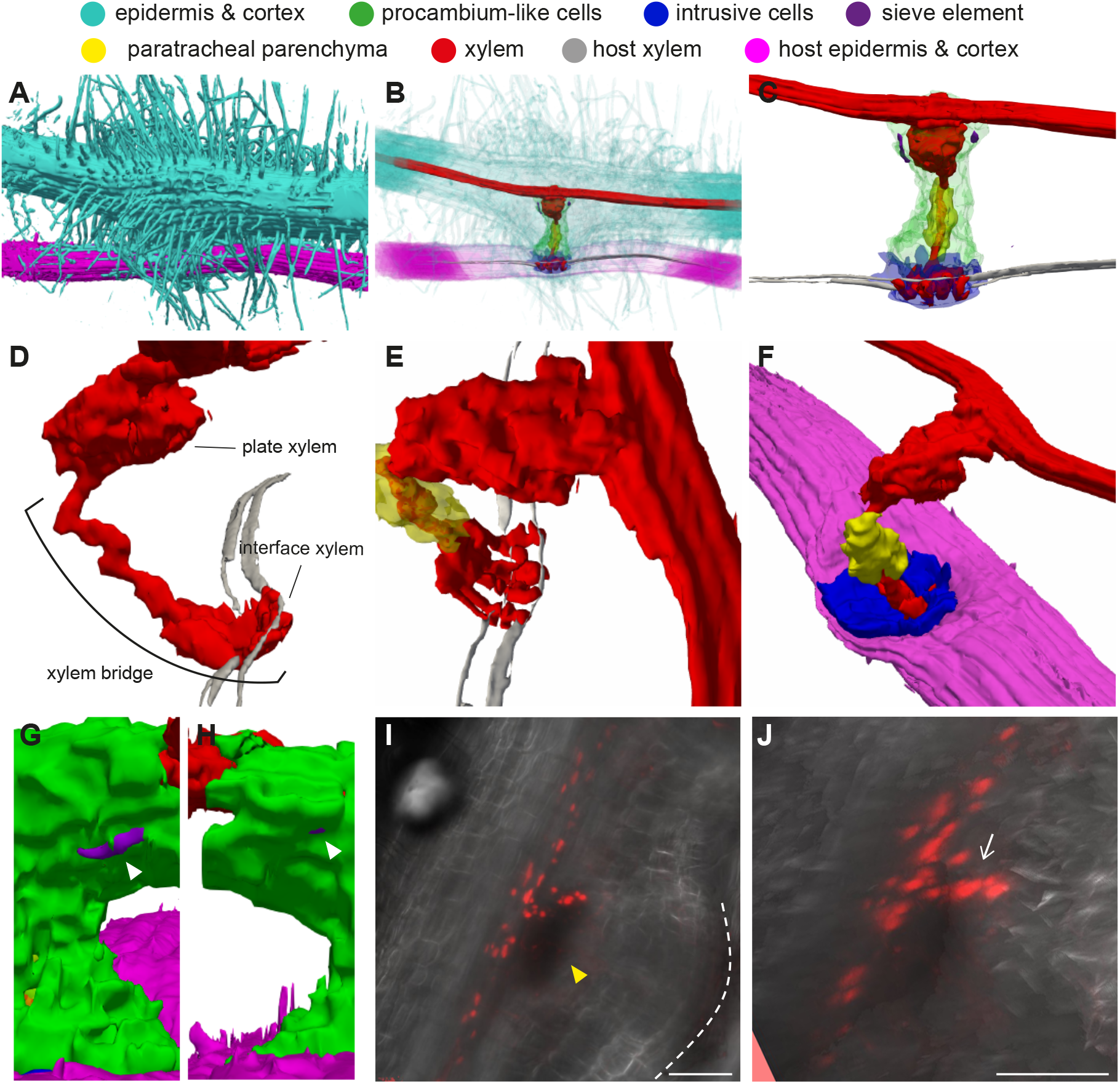
3-D images of a *P. japonicum* haustorium. A, Surface view showing attachment of many haustorial hairs on a host root. B, Semi-transparent view of a haustorium and the host root. C, Internal haustorial cells and host root vasculature. D, E, Close-up views of a xylem bridge interacting with the host’s xylem. F, Close-up view of intrusive cells that cover the entire interaction surface with the host. G, H, Close-up views of sieve elements (white arrowheads) from both sides. I, J, Stacked confocal microscopy image of *pAtPEAR1::tdTomato-nls* expression (red signals) 14 days after infection. J shows the magnified image of I from a different angle, showing the red signals are branching from the main vasculature of a *P. japonicum* root. The plate xylem is located in the dark region indicated by a yellow arrowhead. White dashed lines indicate the outline of the haustorium. Bars = 100 μm.

## Discussion

In this study, we examined the spatial arrangement of cell types within the haustoria of two closely related Orobanchaceae species, *S. hermonthica* and *P. japonicum*. Three-dimensional visualization of haustoria combined with detailed intra-cellular observations revealed the characteristics of each cell type and differences between the parasitic species.

### Hyaline body and paratracheal parenchyma cells

The most prominent difference in the haustorial structure between the two species of parasitic plants was the presence or absence of a hyaline body. The hyaline body in *S. hermonthica* infecting rice forms a ring-shaped thick layer surrounding the procambium-like cells and the xylem bridge. In contrast to *S. hermonthica*, we did not observe such a hyaline body in *P. japonicum*, in which there were no cells with large intercellular spaces accumulating extracellular deposits or with organelle-rich dense cytoplasm. Instead, we observed PP cells that were located along a xylem bridge and had dense cytoplasm and thick primary cell walls at the interface with xylem elements. Similar cells have been reported to be present in other facultative parasites, *Rhinanthus minor*, *Odonites vernus*, *Melampyrum pretense* (Pielach et al., 2014) and *Triphysaria versicolor* (Heidejorgensen and Kuijt, 1995). Similar terms, such as plasma-rich parenchyma cells in the obligate parasite *Orobanche crenata* (Dörr and Kollmann, 1976) and paratracheal parenchyma in *S. hermonthica,* have also been reported (Olivier et al., 1991); however, the parenchyma cells in *O. crenata* or *S. hermonthica* were located rather closer to or adjacent to host xylem vessels than the parasite xylem bridge. Therefore, the functions of these PP cells in facultative parasites and those described in obligate parasites may differ. In the *S. hermonthica* and rice interaction, we did not find any PP-like cells with thickened cell walls next to the xylem bridge (Fig. 2C). PP cells have been proposed to transfer xylem materials to the hyaline body where xylem-derived materials are transiently stored or processed (Pielach et al., 2014). According to this hypothesis, the function of *P. japonicum* PP cells remains obscure in the absence of an apparent hyaline body. Hyaline body formation relies on host compatibility, as shown in the facultative *R. minor* infecting the incompatible host *Prunella vulgaris* (Pielach et al., 2014). Notably, the incompatible interaction of *R. minor* is not able to develop a xylem bridge. Although growth of *P. japonicum* is benefited from *Arabidopsis* infection (Spallek et al., 2017), it is possible that *Arabidopsis* is not sufficiently compatible with *P. japonicum* for hyaline body formation. Alternatively, the lack of a hyaline body may be a natural feature of *P. japonicum* haustoria.

### Xylem connection

The structure of xylem bridges, one of the most conserved features among parasitic plant haustoria, was markedly different between *S. hermonthica* and *P. japonicum,* particularly at the junction points with the host xylem. In *S. hermonthica*, the xylem bridge was perpendicular to the host root, relatively straight and the interfacing xylem cells made an extreme turn at the tip in the direction parallel to the host root axis (Fig. 4E). This structure probably resulted from the intrusive cells that first inserted between the host xylem precursor cells and later synchronously or successively differentiated into xylem cells with the neighboring host cells. In addition, some intrusive cells of *S. hermonthica* can directly penetrate the xylem vessels of *Zea mays* and *Sorghum bicolor* in a small area of their cell tips by forming oscula (Dorr, 1997). The oscula structures were also observed in our experiment with rice (Fig. 2L). Differentiation of oscula-forming intrusive cells into vessel elements can eventually lead to xylem continuity (Fig. 3A), providing an oscula-mediated water conducting channel between the host and parasite. Similar connecting structures have been reported in the facultative parasite *Castilleja* (Dobbins and Kuijt, 1973).

Compared to *S. hermonthica*, the xylem bridge of *P. japonicum* is arc-shaped with the interfacing xylem curving around the host xylem to eventually target the host xylem in the distal region (Fig. 6E). Such morphology of the interfacing xylem represents the invading direction of intrusive cells because the interfacing xylem is derived from some intrusive cells following the completion of penetration (Fig. 3B). Given that the *Arabidopsis* root is thin, rapid polar growth of intrusive cells might fail to capture the position of the host’s central vasculature in the proximal region, and, therefore, growth around the host xylem may be due to guidance toward the host’s vasculature. This hypothesis is based on a general assumption that an attraction signal for the tip of intrusive cells may be present in host vasculatures. Such a guidance system is known for pollen tube attraction by synergid cells during pollination, where peptide signals are emitted to attract pollen tubes (Okuda et al., 2009). Interestingly, many haustorial genes are derived from flower genes, suggesting that pollen genes might have been co-opted by the haustorium, as both pollen and haustorial growth involve polar cell growth (Yang et al., 2015; Yoshida et al., 2019). Neither the oscula structures nor the cell wall continuity between the interface xylem and the host xylem was detected at the interface of *P. japonicum* and *Arabidopsis* (Fig. 3B, Fig. S2). Wrapping the interface xylem around the host xylem vessels increases the interacting surface with host cells; therefore, this phenomenon may contribute towards the efficient acquisition of host xylem contents through xylem pits. A similar interaction was reported in the stem parasite *Cuscuta*, in which the searching hyphal tips grasp the host sieve elements with finger-like protrusions (Dawson et al., 1994). Thus, grasping host vascular cells may be a common feature of parasite intrusive cells (e.g., searching hyphae in dodder) that was acquired *via* convergent evolution. Note that the different xylem interacting structures observed in *S. hermonthica* and *P. japonicum* do not reflect differences between obligate and facultative parasites nor terminal and lateral haustoria. Oscula structures were observed in the facultative parasites, *R. minor* and *Castilleja* (Dobbins and Kuijt, 1973; Cameron et al., 2006), whereas the obligate parasite *O. crenata* and the facultative parasite *T. versicolor* do not form oscula, interacting with host xylem via xylem pits (Dörr and Kollmann, 1976; Heidejorgensen and Kuijt, 1993). The interacting structure could also be affected by host species or compatibility with hosts. Observations in different host-parasite combinations is required to formulate general conclusions on the structural characteristics of the xylem connection and its relationship with host dependency.

### Discontinuity of sieve elements between a host and a parasite

Both *S. hermonthica* and *P. japonicum* have SE in the basal region of haustoria in the vicinity of the plate xylem (Fig. 2C, 5L). Our 3-D reconstruction combined with marker expression analysis revealed that these SE are connected to phloem strands emanating from the parasite’s main vasculature but not to those of the host’s roots (Fig. 4D, 6C). These SE are interspaced with host phloem by procambium-like cells. A similar observation was reported in *Alectra* where haustorial and host phloem elements are interspaced by haustorial parenchymatic cells (Dorr et al., 1979). Based on election microscope observations and *pSUC2-GFP* translocation assays, which can visualize intercellular simplistic continuity between host and parasite, phloem connections are thought to occur only in holoparasitic plants (Dorr and Kollmann, 1995; Dorr, 1997; Ekawa and Aoki, 2017; Spallek et al., 2017; Krupp et al., 2019). This observation reflects the importance of phloem-mediated nutrient transfer in holoparasites whose sugar acquisition completely depends on the host. Important questions remain as to how hemiparasites translocate organic compounds with only xylem connections. Tracer experiments with carboxyfluorescein diacetate (CFDA) dye revealed that, although CFDA translocation from the host to the holoparasite *Phelipanche ramose* is through haustorial phloem strands (Peron et al., 2016), transport to *P. japonicum* is through the inner region of haustoria where intrusive cells and procambium-like cells are located (Spallek et al., 2017). Without characteristic SE in the inner haustorium of hemiparasites, other cell types may function in nutrient delivery either in a symplastic or apoplastic manner.

### Other cell types

Our FE-SEM analyses revealed several novel features in *P. japonicum* haustorial cells. Vacuoles of procambium-like cells have frequent membrane invaginations and contain many single- or multi-membrane vesicles (Fig. 5G), which indicates active degradation of cytosolic metabolites and organelles. These vesicles are likely associated with vacuole autophagy, a process that can be induced by nutrient starvation or biotic stress such as pathogen attack (Yu, 1999; Cao et al., 2018; Yoshimoto and Ohsumi, 2018). Whether there is direct functional relevance of the autophagy process in plant parasitism remains to be investigated. In addition, we found two types of procambium-like cells with distinct subcellular features (Fig. 5F). Whereas those in the center and base of the haustorium are highly vacuolated (Fig. 5G), those proximal to the host contain many atypical membrane structures in the cytosol (Fig. 5H). Subtle differences also exist in terms of marker gene expression in the procambium; whereas, Pj*HB15a*, Pj*HB8* and Pj*WOX4* were expressed widely in all procambium-like cells, the expression of Pj*HB15b* was limited to the center and basal regions (Wakatake et al., 2018). The identification of additional cell type-specific marker genes is necessary to verify the identities of haustorial cell types in the future.

## Materials and methods

### Plant materials and growth conditions

The parasitic plants used in this study were *Phtheirospermum japonicum* (Thunb.) Kanitz ecotype Okayama and *Striga hermonthica* (Del.) Benth. and the host plants were *Arabidopsis thaliana* ecotype Columbia (Col-0) and japonica rice (*Oryza sativa* L.) cultivar Koshihikari, respectively. Seed sterilization and host infection were performed as described previously (Cui et al., 2018). Briefly, one-week preconditioned *S. hermonthica* seeds were treated with 10 nM strigol for 1 d and placed onto rice roots that had been grown using a rhizotron system for 1 week after germinating one week on moistened filter paper. The *S. hermonthica*-infected rice plants were grown under long-day conditions (16 h light/8h dark, 90 μmol m^−2^s^−1^) at 25°C for 2 weeks. *P. japonicum* and *Arabidopsis* were germinated on half strength Murashige and Skoog (MS) solid media at 25°C and 22°C, respectively, under long-day conditions. Seven-day-old *P. japonicum* seedlings were transferred to a nutrient-free agar medium and cultivated for 3 d under long-day conditions for the starvation treatment. The main root of 7-day-old *Arabidopsis* seedlings was then placed next to the root of *P. japonicum* to induce infection. One day after infection, successfully induced haustoria were marked; only marked haustoria were collected for sectioning at 9 days after infection. Infection was carried out at 25°C under long-day conditions.

### Preparation of serial semi-thin sections of haustoria

Two-week-old *S. hermonthica* haustoria and 9-day-old *P. japonicum* haustoria were collected by excising the roots of the host and parasite 2~3 mm above and below the infection sites. The haustorial samples were fixed with 4% (w/v) formaldehyde, 2% (v/v) glutaraldehyde and 0.05 M cacodylate buffer, dehydrated in a methanol dilution series, and embedded in EPON 812 resin (TAAB). The resin blocks were sliced with an ultramicrotome (Leica microsystems EM-UC7) using a diamond knife (DIATOME Histo), and 1 μm-thick serial sections were prepared. Serial sections were stained with 0.05% (w/v) toluidine blue, and photographs were taken using an upright light microscope (Olympus BX51M & DP26 digital camera).

### Field emission scanning electron microscopy (FE-SEM) observations

Toluidine blue-stained serial sections, prepared as describe above, were stained with a 0.4% (w/v) uranyl acetate solution for 12 minutes followed by a lead citrate solution for 3 minutes, dried and coated with osmium for 1 second using an osmium coater (HPC-1SW). Observations and imaging were accomplished using a field emission scanning electron microscope (Hitachi High-Tech SU 8220) equipped with a highly sensitive backscatter-electron detector (YAG-BSE 5 kV).

### 3-D reconstruction of surface structures

#### 3-D alignment adjustment

To construct 3-D volumetric images, serial images of toluidine blue-stained sections were adjusted for translation and rotation by rigid image registration using a Python library in SimpleElastix (Marstal et al., 2016). In addition to rigid registration, intensity based non-rigid image registration was applied to iron out wrinkles in the sectioned images. For non-rigid image registration, wrinkled images were registered to their respective nearest wrinkle-free image. After automated alignment adjustment, section alignment was manually checked, and misaligned sections were re-registered by changing the registration parameters. The tools are available at https://github.com/yk-szk/ssrvtools.

#### Semi-automated cell labeling

For color-coding of cell types, we developed a semi-automated labeling method. Each section image was split into RGB channels, and an outline was extracted using the R channel or G channel images using line enhancement and removal of background noise. After line extraction, each line-enclosed area was defined as a single cell. Every fifth slice was subjected to manual painting, in which different cell types within the haustorium were marked manually with different colors using Photoshop CS6 (Adobe) and a tablet pen. We developed a Python script (https://github.com/yk-szk/ssrvtools) that automatically paints the intercepted unpainted slices based on information from manually painted images and single cell outlines. The manually painted images and single cell outlines of unpainted images were over-laid, and the color occupying the largest area in the cell was selected to paint the cell. This procedure was repeated up to four times based on one manually painted image. After automatic painting, manual corrections were applied whenever necessary.

#### Polygon mesh extraction and visualization

Color-painted images were stacked to construct a color-coded 3-D volume image. The surface structure of each cell type was extracted as a polygon mesh using Marching Cubes (Lorensen and Cline, 1987). Extracted polygon meshes were visualized using the open-source visualization application Paraview (Kitware).

### Plasmid construction and transformation

The *Arabidopsis PEAR1* (At*PEAR1*) promoter sequence was amplified by the primers (5’-TCGACTCTAGAGGATCCCCGGGTGTTGCCTAACTCTTGATTATTGATT-3’ and 5’-CCCTTGCTCACCATCCCTGGTTATTCTCTTTTGATTTTATTCTTCAAAAT-3’) and cloned into the pAN19 vector by the SLiCE technique (Okegawa and Motohashi, 2015) using the coding sequence of the *tdTomato-NLS* reporter. The *p*At*PEAR1::tdTomato-NLS* sequence was inserted into the modified pBIN19 binary vector containing the Basta herbicide resistance gene (Miyashima et al., 2011). The binary vector was transformed into *P. japonicum* seedlings as previously described (Ishida et al., 2016). The binary vector was transformed into *Agrobacterium rhizogenes* strain AR1193. Five-day-old *P. japonicum* seedlings were suspended in the *Agrobacterium* solution, sonicated and vacuum infiltrated. The infiltrated seedlings were cultured on Gamborg’s B5 medium with 0.5% (w/v) agar, 1% (w/v) sucrose and 450 μM acetosyringone (Sigma) at 23°C in the dark for 2 days. The seedlings were transferred to Gamborg’s B5 medium with 0.5% (w/v) agar, 1% (w/v) sucrose and 300 μg/ml cefotaxime (Tokyo Chemical Industry), and then cultured at 25°C under long-day conditions (18h/6h for light/dark, 90 μmol m^−2^s^−1^) for 18 days, to initiate hairy roots. Initiated hairy roots expressing the fluorescence signal were cultured on the same medium for 2 weeks, and then transferred onto 0.8% (w/v) agar plates for the starvation treatment. After a 1-day starvation treatment, the hairy roots were used to infect *Arabidopsis* roots. A 2-week old haustorium expressing a fluorescence signal was observed by Nikon A1 confocal microscopy.

### Confocal microscope observation

For xylem staining, haustorial and host root segments were excised from the roots, stained by SR2200 (Renaissance) and Fuchsine, and cleared by ClearSee solution by following previously described protocols with a slight modification (Ursache et al., 2018; Miyashima et al., 2019). The haustorial samples were submerged in 0.1% (v/v) SCRI Renaissance 22000 solution in 1×PBS buffer, vacuum infiltrated for 15 min and incubated at 4°C for three days. The sample was then washed once with 1× PBS buffer, immersed in 500 μl of ClearSee solution (10% (w/v) xylitol, 15% (w/v) deoxycholate, 25% (w/v) urea) supplemented with 140 μl of 0.2% (v/v) Fuchsine, and placed under vacuum until no bubbles leaked out from the sample. After repeating this step twice, the sample was left at room temperature for 4 h. The solution was then replaced by new ClearSee solution, subjected to a short vacuum treatment and incubated overnight at room temperature. For imaging, a Zeiss LSM710 confocal microscope was used with excitation and emission wavelengths suitable for blue and red colors.

## Acknowledgement

We thank Dr. A. G. T. Babiker (University of Khartoum, Sudan) for providing *S. hermonthica* seeds, Dr. Kenji Mori for providing strigol. We also thank Ms. Kana Tsurii (RIKEN) and Dr. Kei Hashimoto (RIKEN) for technical assistance in sample preparation and FE-SEM observation.

## Supplemental Data

**Supplemental Figure S1.** Adjustment of coloring borders during automatic interpolation. Cell type-specific colors derived from the original slide results in the border shift in the next slide (left panels) due to changes in cell volume between each section. This phenomenon causes the appearance of multiple colors in a single cell at the cell type-specific border (an example is shown by an asterisk in the top left panel). Border adjustment is applied by conferring the cell with the color that originally occupied the majority of the cell area (asterisk in the top right panel). In the case of cells at the organ boundary (bottom panels), color borders are adjusted to be fully contained inside the cells.

**Supplemental Figure S2.** Xylem-xylem interaction between *P. japonicum* and *Arabidopsis*. Confocal microscope images of a *P. japonicum* haustorium infecting an *Arabidopsis* root. The haustorium was stained by SR2200 (Renaissance) for cell outline (blue) and Fuchsine (red) for xylem. A, A stacked image of a haustorium from serial projections. *Arabidopsis* root xylem cells are oriented laterally and *P. japonicum* interface xylems are aligned perpendicularly. B, A single projection image from A. C, A lateral view of a haustorium at the plane indicated by a dashed line in B. Note that the cell walls of the interface xylem (arrowhead) and host metaxylem (arrow) remain intact.

**Movie 1.** An opaque 3-D structure of an *S. hermonthica* haustorium-infected rice.

**Movie 2.** A semi-transparent 3-D structure of an *S. hermonthica* haustorium-infected rice.

**Movie 3.** A 3-D internal structure of an *S. hermonthica* haustorium-infected rice.

**Movie 4.** An opaque 3-D structure of a *P. japonicum* haustorium-infected *Arabidopsis.*

**Movie 5.** A semi-transparent 3-D structure of a *P. japonicum* haustorium-infected *Arabidopsis*.

**Movie 6.** A 3-D internal structure of a *P. japonicum* haustorium-infected *Arabidopsis.*

